# The duplication of genomes and gene regulatory networks and its potential for evolutionary adaptation and survival

**DOI:** 10.1101/2023.04.28.538696

**Authors:** Mehrshad Ebadi, Quinten Bafort, Eshchar Mizrachi, Pieter Audenaert, Pieter Simoens, Marc Van Montagu, Dries Bonte, Yves Van de Peer

## Abstract

The importance of whole genome duplication (WGD), or polyploidy, for evolution, is controversial. Whereas some view WGD mainly as detrimental and an evolutionary dead end, there is growing evidence that (the establishment of) polyploidy can help overcome environmental change, stressful conditions, or periods of extinction. However, despite much research, the mechanistic underpinnings of why and how polyploids might be able to outcompete or outlive non-polyploids at times of environmental upheaval remain elusive, especially for autopolyploids, in which heterosis effects are limited. On the longer term, WGD might increase both mutational and environmental robustness due to redundancy and increased genetic variation, but on the short – or even immediate – term, selective advantages of WGDs are harder to explain. Here, by duplicating artificially generated Gene Regulatory Networks (GRNs), we show that duplicated GRNs – and thus duplicated genomes – show higher signal output variation than non-duplicated GRNs. This increased variation leads to niche expansion and can provide polyploid populations with substantial advantages to survive environmental turmoil. In contrast, under stable environments, GRNs might be maladaptive to changes, a phenomenon that is exacerbated in duplicated GRNs. We believe that these results provide new insights into how genome duplication and (auto)polyploidy might help organisms to adapt quickly to novel conditions and to survive ecological uproar or even cataclysmic events.

**Statement of Relevance:** In everyday life, it is often critical to be able to evaluate the quality of our own cognitive decisions and actions. However, one of our most frequent actions often does not even reach our awareness: eye movements. We investigated whether observers were able to successfully judge the accuracy of their eye movements when tracking a cloud of dots that followed an unpredictable trajectory. We found that observers were able to distinguish good from bad trials, although sensitivity was lower compared to equivalent previous reports when judging the quality of hand movements. These results add an 35 item to the growing list of our metacognitive abilities, but critically for eye movements that we are typically unaware of. They lead to important novel questions about why metacognitive abilities differ across decisions or different types of actions, and what the potential components of metacognitive
abilities might be.

## Introduction

Whole-genome duplication (WGD) leading to polyploidy is a common phenomenon that has been studied for over 100 years, especially in flowering plants (Soltis et al. 2014). Because of the well-known detrimental effects arising from genome doubling, most WGD events are not successful. Genomic instability, mitotic and meiotic abnormalities, and minority cytotype exclusion are all expected to quickly remove new polyploids from the population (Levin 1975; Comai 2005; Morgan et al. 2020). Nevertheless, there are numerous polyploid organisms around us. Furthermore, even those organisms that are currently considered ‘functional’ diploids usually bear signatures of a polyploid ancestry (Wendel 2015; Van de Peer et al. 2017). Several of these ancestral polyploidy events can be traced back to the origin and diversification of major phylogenetic lineages, including vertebrates, fishes, and flowering plants; and within flowering plants, core eudicots, monocots, orchids, grasses, composites, and legumes (Taylor et al. 2003; Dehal and Boore 2005; Van de Peer et al. 2017).

This phylogenetic signal of polyploidy success suggests an important role for WGD in promoting phenotypic diversity, with a subsequent facilitating role in speciation (Soltis and Soltis 2009; Landis et al. 2017; Leebens-Mack et al. 2019). Speciation typically occurs under restricted conditions where certain genotypes can exploit novel ecological opportunities under the presence of mating barriers with others (Schluter 2001). More importantly, polyploidisation is often associated with the expression of new, often exaggerated, phenotypes that have the potential to promote niche expansion and a subsequent radiation in novel environments. Doubling the amount of DNA does for instance necessitate larger cell nuclei and cell size and has already major consequences on organismal developmental and physiological responses (Baduel et al. 2018; Bomblies 2020). Size independent phenotypic changes have been documented on stress physiology and other traits that provide advantages under extreme environments (te Beest et al. 2012; Van de Peer et al. 2021). It has been suggested that such potential niche expansion advantages in novel environments might be responsible for phylogenetic records showing a rise of polyploids at certain epoch boundaries, such as for instance the K-Pg boundary, a geological period characterized by major episodes of global climatic change and mass extinction (Fawcett et al. 2009; Vanneste et al. 2014a; Van de Peer et al. 2017; Cai et al. 2019; Koenen et al. 2020; Wu et al. 2020; Chang et al. 2023), or around recent glaciation maxima (Novikova et al. 2018). Studies in yeast have shown that polyploidy can accelerate evolutionary adaptation to challenging environments, because WGD induced regulatory redundancy followed by divergence, allowing a wider range of phenotypic responses to environmental stresses (Selmecki et al. 2015; Scott et al. 2017).

The increasing numbers of genes that diversify in function due to a relaxed functional constraint on one of both copies (i.e., sub- or neofunctionalization (Force et al. 1999)), is likely not the sole explanation of WGD’s evolutionary success under stress. Doubling of gene regulatory networks may equally increase the frequency of beneficial mutations (Selmecki et al. 2015), and therefore enlarge the genetic and phenotypic variation for selection to act on. In this respect, WGD can be seen as a complex super mutation of/within the genome. The increased genetic variation and the buffering effect of their duplicated genes has led to an increased recognition of the adaptive potential of polyploidy (Van de Peer et al. 2009; Van de Peer et al. 2017; Doyle and Coate 2019).

Recent work by some of us based on a computational framework aimed at mimicking biological evolution (Yao et al. 2014; Yao et al. 2016) suggested that so-called digital organisms (DOs) with a ‘simple’, unduplicated genome performed better - as in, adapted faster - than DOs with a duplicated genome in stable environments, while the opposite was true for unstable environments (Yao et al. 2019). Somewhat similar observations were made by Cuypers and Hogeweg (Cuypers and Hogeweg 2014) and Cuypers et al. (Cuypers et al. 2017) using populations of so-called ‘virtual cells’. These insights were generated by the implementation of WGD as a series of random mutations of large – adaptive or maladaptive – effect. Gene regulatory networks (GRNs) shape the mechanistic pathway between genotype and phenotype. We build on the observation that duplicated GRNs seem to have magnified impact and hypothesize that the phenotypic variance generated by such networks exceeds the one of the ancestral simpler networks. The eventual propagation of information through (artificial) networks, and thus the eventual distribution of output signals is uncertain. Signal propagation across a network can be considered as a sum of different node values. The eventual variance of the distribution of output signals in such systems with a double number of nodes will then be the sum of the variances of the distribution of output signals across all nodes and their (doubled) covariance. Hence, increasing the number of nodes, and given covariances not being strongly negative, variance of the distribution of output signals of the population of doubled networks should always increase. To what extent the duplicated structure of the network results in a different signal propagation relative to the non-duplicated version, or to networks with the same number of nodes but with random structure, remains understudied. We developed simulations with artificial gene regulatory networks (aGRNs) of different sizes and shapes and studied the potential consequences of environmental change for diploid and polyploid organisms.

## Methods

### Defining and initializing aGRNs

In the current study, we consider artificial gene regulatory networks (aGRNs), mimicking gene regulatory networks in the traditional sense i.e., a set of genes or proteins that interact with each other to define and control a specific function. For instance, such networks can transduce signals from environmental cues into a proper phenotypic behaviour that allows an organism to respond to environmental changes. In our aGRNS, we discriminate between ‘regulatory’ genes or proteins (like transcription factors, TFs), regulating the activity of other genes or proteins, and so-called ‘output’ genes or proteins, which produce an ‘output’, such as a structural protein or a metabolite. We also consider so-called ‘input’ genes or proteins, which can ‘sense’ the environment, and which can receive an input value. All these different genes or proteins form the nodes of the network, while edges between nodes represent their interactions. Furthermore, the following rules apply: 1) networks have a fixed number of nodes and are built by a preferential attachment algorithm and thus have properties that are close to scale-free networks, and 2) all edges are directed and have weights to mimic the strength of regulation (interaction). For instance, a weight can be considered as the strength with which a regulator binds to its target, or alternatively, as the strength with which it induces - or represses - expression of its target. It should be mentioned that all ‘simple’ or ‘single’ (non-duplicated, ancestral) networks have two output nodes (while the duplicated network has four output nodes). As a result, for better interpretation of the outcomes, plots are two-dimensional (see below).

Although there is still debate as to what extent biological networks are truly ‘scale-free’ (Broido and Clauset 2019), there are reasons to believe that many biological networks at least have certain features similar to scale-free networks, such as a high diversity of node degrees and absence of nodes in the network that could be used to characterize the rest of the nodes (Albert 2005). Therefore, here, we consider directed, weighted, scale-free networks as our initial networks. To generate these directed scale-free networks, we used the ‘Preferential Attachment’ algorithm (Barabasi and Bonabeau 2003; Barabasi 2009; Prettejohn et al. 2011). Using this algorithm, nodes have a higher chance to connect with nodes with a higher degree (more connections) compared to other nodes (‘rich get richer’) (Barabasi and Bonabeau 2003). Another significant characteristic of real biological GRNs is the high number of feed-forward loops (FFL; A regulates B, B regulates C, A regulates C)(Cooper et al. 2008; Mottes et al. 2021). To enrich our aGRNs with FFLs, we used the algorithm of Herrera and Zufiria (Herrera and Zufiria 2011). By using this algorithm, the clustering coefficient of the network increases which in turn causes an increase in the number of triadic motifs in the network. Then, by controlling and changing the direction of edges, we can easily raise the number of FFL motifs in the network. It should also be noted that, for computational reasons, generated networks containing feedback loops (never-ending loops; A regulates B, B regulates C, and C regulates A, or self-regulation like A regulating A) are discarded. Besides, although such motifs do occur in real biological networks, they are rare (Tran et al. 2013).

As stated above, in our aGRNs, three different kinds of nodes are distinguished (Fig. 1). Nodes with zero in-degree and non-zero out-degree are referred to as input nodes. The number of input nodes is variable and depends on the preferential attachment algorithm, but usually lies between 20% and 40%. Nodes with non-zero in-degree and non-zero out-degree represent regulators, affecting other nodes (genes), such as transcription factors. Finally, nodes with non-zero in-degree and zero out-degree are output nodes defining the ‘phenotype’ or ‘behaviour’. To allow analyses of the phenotypic effects in an easily conceivable two-dimensional space (see further), the number of output nodes is artificially set to 2. In practice, this means that, for a network with n nodes, the network is built on n-2 nodes, and after the n-2 network has been built, the last two nodes are added. These two nodes cannot connect to others, but others can connect to them, again by applying the preferential attachment algorithm. Every edge has a weight corresponding to the ‘strength’ of the regulatory interaction between genes. The weight of the edges is determined randomly by a standard Normal Distribution, generating values between -1 and 1, with positive values indicating stimulation, and negative values repression. After initialization of the network, the weights of the edges are fixed. However, to model changes in the network after receiving different values of input nodes, i.e., mimicking environmental cues, we have defined an ‘activation level’ for each node. Initially, activation levels of all nodes are set to 0, but during simulation, the input signal will determine the activation level of the input node(s), which will then be propagated through the network changing the activation level of each downstream node in function of all incoming edge weights and the activation level of all previous nodes (Moreira and Rennó-Costa 2021). Concretely, this way, when the value/expression of one node/gene is increased (or decreased), this would lead to increased (or decreased) dosage of a regulator, in turn being responsible for the increased (or decreased) production of its target, and so on (see below).

**Fig. 1.**
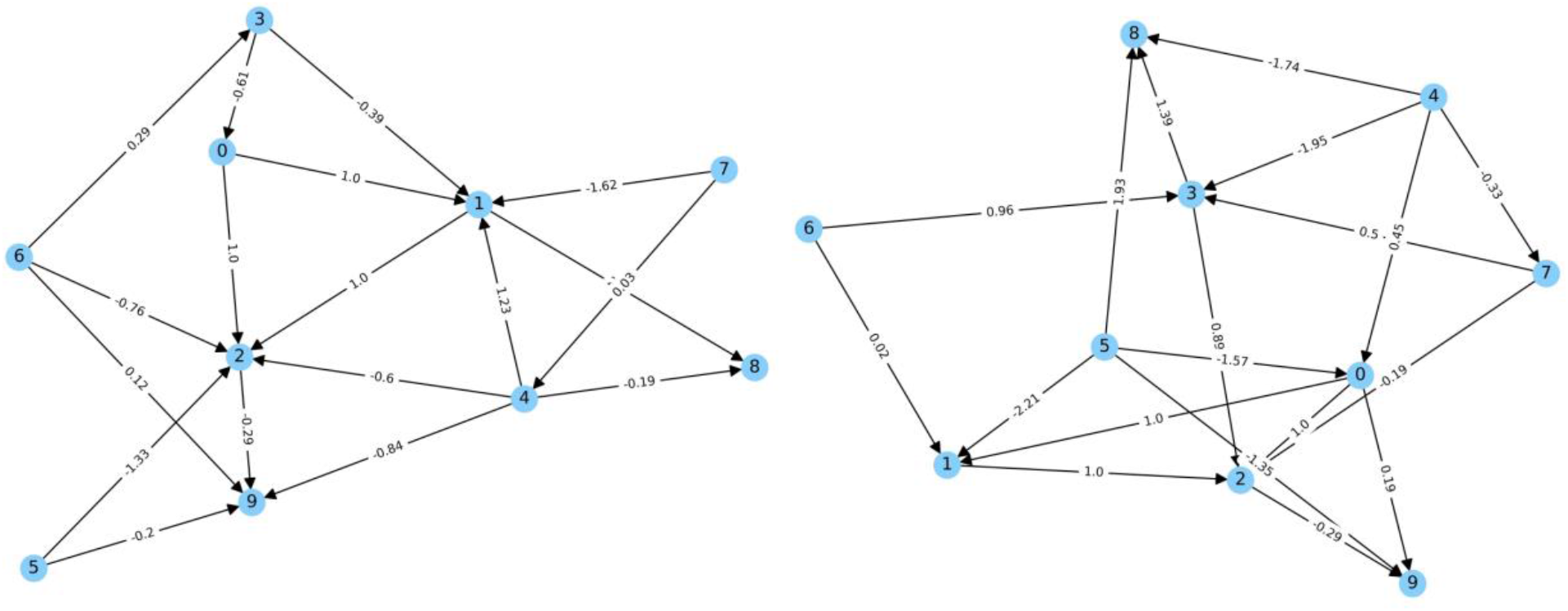
Two examples of an aGRN of 10 nodes generated by the preferential attachment algorithm. All nodes represent regulatory genes or proteins, except nodes 8 and 9 in both networks, which are output nodes. Nodes 5 and 6 can act as input nodes since all edges are outgoing. Weight values are also indicated. Positive weight values represent induction, while negative weight values indicate repression (as for example in gene expression). The topology of a specific aGRN is unique and can be considered the genotype, while the output nodes or node values define the phenotype. See text for details.

### Network duplication

To mimic WGDs, we simply duplicate all nodes of the network. However, this means that, if a regulator A regulates nodes B and C, its duplicate A’ regulates the duplicated targets B’ and C’, but also the original targets B and C. In turn, the original regulator A also regulates all four targets, B, B’, C, and C’ (Fig. 2). The edge weights between corresponding nodes in the non-duplicated (A-B) and duplicated (A-B, A’-B’, A’-B, A-B’) network remain unaltered. It should be noted that such operation mimics only part of the effects of a polyploidisation event, i.e., the effects of genome doubling (autopolyploidy) and not those of genome merging (allopolyploidy). As a result, throughout the paper, we will only consider autopolyploidy, where the ‘own’ genome gets duplicated, rather than allopolyploidy, where the duplicated genome is obtained from the merging of genomes of different species. Although there is still discussion on the ratio of autopolyploids versus allopolyploids in the polyploid realm, there is reason to believe that autopolyploids are much more frequent than previously thought (Barker et al. 2016). We are aware of the fact that, in autopolyploids, unlike in allopolyploids, the duplicated genes might be seen more as different alleles of the same gene, rather than as different genes, but we feel this will not have a major effect on our conclusions, because even when considered only different alleles, it will affect certain traits (e.g., due to dosage effects) (Baduel et al. 2018; Bomblies 2020; Van de Peer et al. 2021), and when there is no recombination, they can be considered separate genes.

**Fig. 2.**
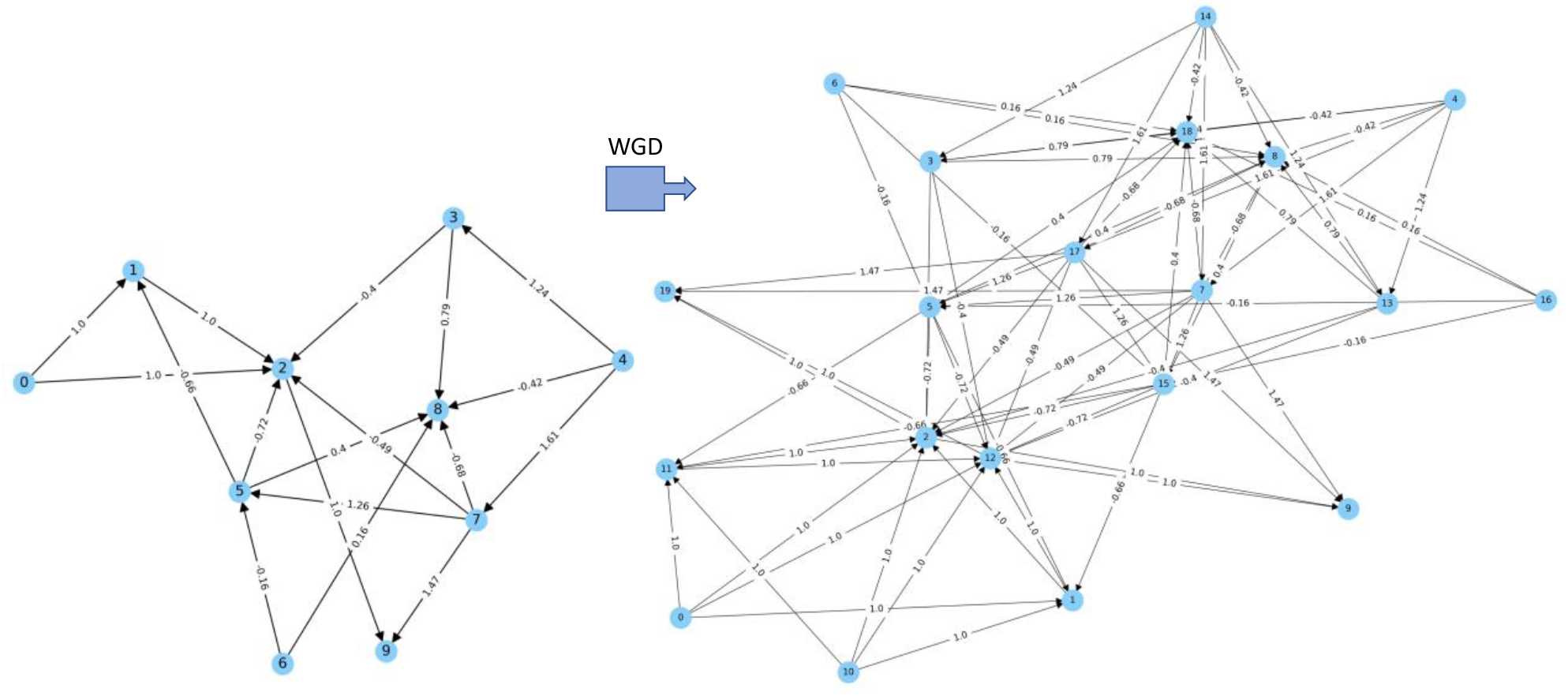
Example of a simple or ancestral (left) and duplicated (right) aGRN.

### Signal propagation in the network

One of the main purposes of our simulations is to see how signals, such as environmental cues, propagate over simple (non-duplicated) versus duplicated networks, the hypothesis here being that, because of the specific structure and a denser wiring of duplicated networks (Fig. 2), greater parts of the network – and thus more genes - are affected, with consequently, greater variation in output values. As far as we know, this has not been studied in a biological context, and certainly not in the context of duplicated networks and polyploidy or genome duplication. Signal propagation functions in the network (see further) will determine the output values, and thus the phenotype. We evaluated output changes by using constrained propagation using the hyperbolic tangent function (tanh) with the max value of “+1” and min value of “-1” (eq. 1). This function is typically used to determine the activation level of nodes in neural networks (LeCun et al. 2015; Nwankpa et al. 2018; Moreira and Rennó-Costa 2021).

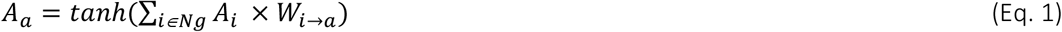

where *A*_*a*_ is the activation level of node *a, W*_*i*→*a*_ is the weight of the edge from node *i* to node *a*, and *Ng* is the list of nodes that are connected to node *A*.

Depending on the different input signals, different outputs will be reached. This constrained implementation mimics biological networks in that it considers minimal and maximum values for, for instance, an increase in gene expression (increase in gene expression is not unlimited). We additionally provide a sensitivity analysis for an unconstrained linear propagation algorithm in *SI Appendix*, Text S1.

Environmental changes, such as for instance changes in temperature, are mimicked by changing the signal values of input nodes (input values are drawn from a uniform distribution between -1 and +1) and following their propagation over the network. Values from the two output nodes, *o1, o2*, are interpreted as a phenotype in a two-dimensional (2D) trait space (hence the two output nodes). Since we have four output nodes for duplicated networks, like *o1, o2, and o1’ and o2’*, each output value is calculated as the average of the corresponding output nodes, e.g. *(o1+o1’)/2*. We thus consider a conservative but realistic full dosage compensation of the gene expression after WGD (Doyle and Coate 2019). This representation allows us to quantify phenotypic changes by means of Phenotypic Trajectory Analysis, PTA (Adams and Collyer 2009; De Lisle and Bolnick 2020). In brief, this approach allows us to understand whether evolutionary divergence between pairs of populations, here the ancestral (single/simple) and duplicated networks, is parallel, convergent, divergent, or random (Bolnick et al. 2018). To this end, vectors 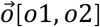 [*o*1, *o*2] are drawn from the ‘phenotype/trait value’ (e.g., positions of the two output nodes in a 2D space, see Fig. *3A*) of an ancestral simple network to the ‘phenotype/trait value’ of its duplicated counterpart. Changes in the vector length demonstrate how much the eventual trait value (the phenotype produced by the duplicated genome) is changing compared to the initial value of the simple aGRN. The distribution of these differential vector lengths therefore identifies the strength of the phenotypic (and putative niche) shifts, and thus the strength of the divergence due to genome doubling. The orientation of a vector in trait-space describes the relative contributions of the output traits to divergence between that pair of populations. Changes in angular dispersion between the ancestral and duplicated phenotype – following changing environmental conditions mimicked by changing the input signal - indicate whether phenotypic changes among all independent network doubling events occur in parallel for all doubling events (absolute angles in the trait space similar, hence showing directional or parallel evolution), in the same direction of the initial phenotypic position (relative angles between the ancestral and doubles genotypes are zero, showing niche expansion), or completely random (both absolute and relative angles randomly distributed across trait space). We calculated overall phenotypic variation at the population-level (hence a population of 10K single versus doubled genomes generated by the same initialization) by calculating the variance of the mean of the two output nodes, and by multiplying the variance of the two output values. Artificial gene regulatory networks consisting of 10, 20, 40, 60, 80, and 100 nodes were generated and exposed to 10,000 environmental conditions by drawing the input values (activation levels for the input nodes) from a uniform distribution between -1, and +1, as stated previously. We report these variances for 400 populations of 10K networks per category (e.g., number of nodes in the single network; Fig. 3*B*). Unless specifically defined, we present data for ancestral (simple) networks consisting of 10 nodes and node/edge values initiated from the uniform distribution. We tested the robustness of our analysis by sensitivity analyses for the full range of node numbers (*SI Appendix*, Text S2) and network initialisation from Gaussian N(0,1) and mixed uniform U[-1,1]-Gaussian N(0,1) distributions (see *SI Appendix*, Text S3).

**Fig. 3.**
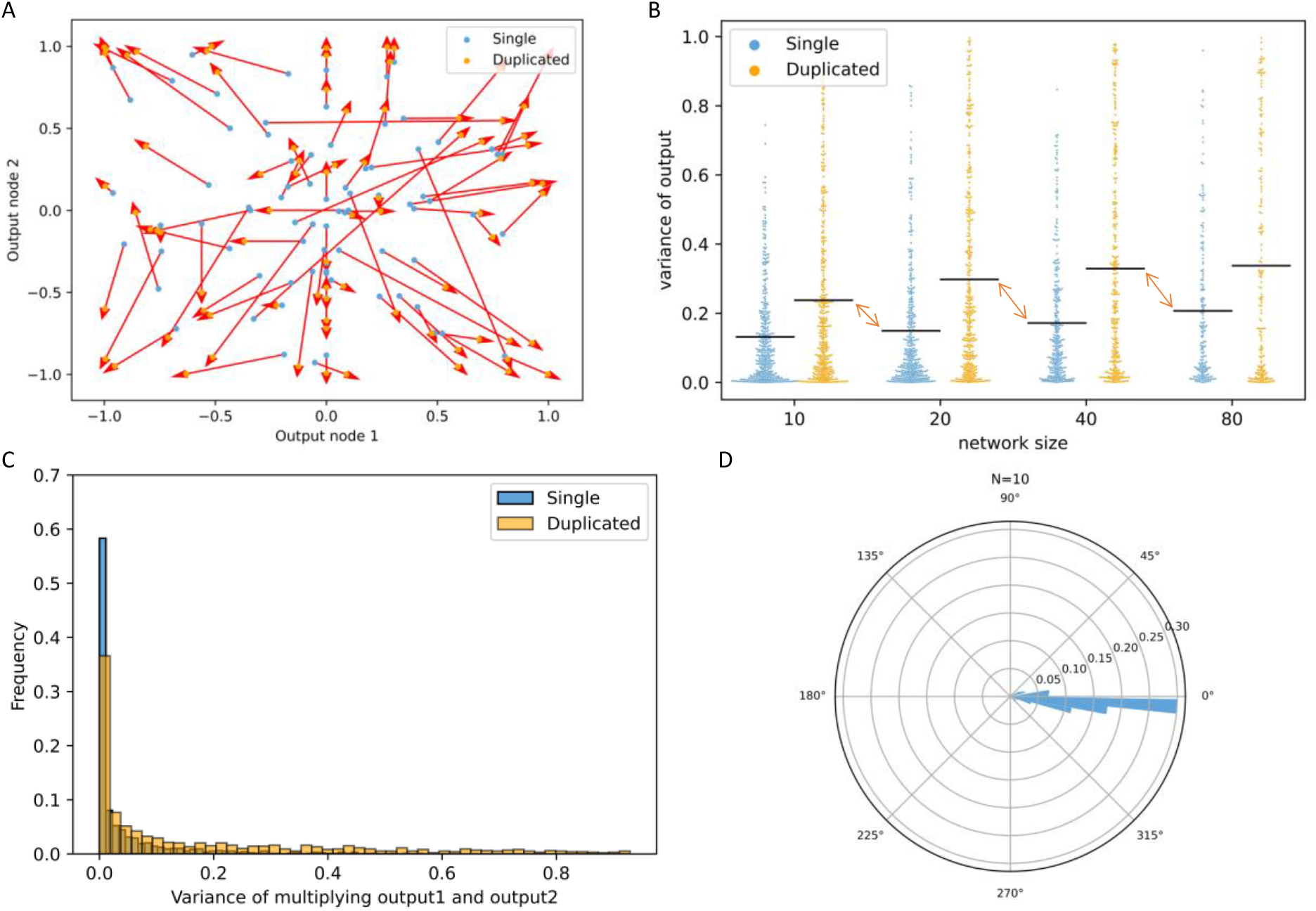
Phenotypic Trajectory Analysis (see Methods) comparing a population of single versus its duplicated networks. (A) The value of one output node is plotted against the value of the second output node for simple (blue dots) and duplicated (orange) networks for a simple GRN of 10 nodes. Thinning has been applied and from the 1,000,000 values only a fraction is shown, to facilitate interpretation. (B) The variance for simple and duplicated networks for networks of 10, 20, 40, and 80 nodes (400 networks consisting of 10K single/double networks per size category). Variance of the output is increasing with node additions, but duplicated genomes always have higher variance compared to their unduplicated counterparts. Red arrows denote the difference in variance between duplicated networks and random networks with an equal number of nodes but not having the typical duplicated topology (structure doubling versus node doubling) (C) Frequency distribution of the phenotypic variance σ as measured by multiplying variance of both (mean) output node values in the 10K simulated simple GRNs of 10 nodes, and their duplicated counterparts. (D) Angular dispersion of the relative angles between the single and doubled networks for 10K simulations of simple GRNs of 10 nodes, and their duplicated counterparts.

Linking phenotypes to the environment: fitness. Besides studying theoretical phenotype-WGD mapping, we moved one step further and tested the hypothesis that duplicated genomes provide fitness benefits under larger environmental changes. To this end, we simplified the two-dimensional phenotype vector 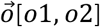[*o*1, *o*2] towards its one-dimensional average value [*ō* = (*o*1 + *o*2)/2] as it was shown to generate qualitatively similar insights (Pearson correlation *r* between 0.45-0.52). We start from a population of simple networks that are well adapted to their environment and assume that the individual phenotypes are all centred around the fitness optimum. We simulated a genome doubling effect of all simple genetic networks and assessed the fitness ***w*** of both the simple and duplicated populations in the reference environment and under environmental changes of different magnitudes. For each size of simple networks (resp. 10, 20, 40, 60, 80, and 100 nodes), we constructed 10,000 simple and their duplicated networks as described above. All these 10K networks have different topologies and different edge weights. For all these simple and duplicated networks, we here provide insights from simulations with an input value of 0.01, creating a large compilation of networks with different output values centered around 0 (see above). This specific input value represents our ‘reference environment’ and guarantees that the average phenotype of the population is close to the fitness optimum (0.01), when we assume that fitness ***w*** is inversely proportional to the difference between input and output value. To assess fitness in the reference environment and how it is affected by the underlying fitness function, the performance of each network was calculated using both a negative linear and a Gaussian fitness function.

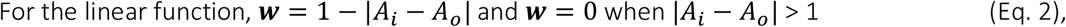

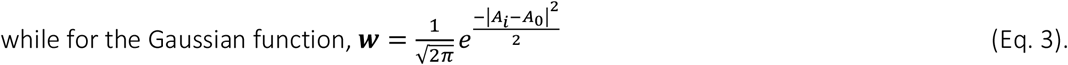

Networks with a phenotype (output *A*_*o*_) similar to the reference environment (input *A*_*i*_) will have the maximal fitness ***w*** = 1, and this value decreases to zero under large deviation from the reference environment.

Next, the populations of simple and duplicated aGRNs were subjected to deviating environmental conditions as input A_i_, our environmental change gradient, and ***w*** was again calculated according to the output phenotype *A*_*o*_ as in Eq. 2 and Eq. 3. To this end, the input value *A*_*i*_ of one randomly chosen input node is changed gradually with *ΔA*_*i*_. If there is more than one input node, which is always the case for duplicated networks, the input of the other input nodes is kept at zero.

### Data availability

Documentation and software to generate artificial scale-free gene regulatory networks and their duplicated versions simulating the result of whole genome duplication (WGD) can be found at: https://github.com/Mehrshad-Ebadi/SC-as-aGRNs. Examples of networks and their duplicated versions of different sizes can be found there as well.

## Results

### The increased phenotypic variance of network duplication

Figure 3*A* shows a Phenotypic Trajectory Analysis (see Methods) of a population of simple (10 nodes) and duplicated networks in a single environment. It provides an example of the increase in the phenotypic variation of the duplicated population compared to the simple (ancestral) ones. Although the phenotypic effect of WGD depends on the network’s topology (i.e., the genotype), the average phenotypic value of the duplicated networks is on average more extreme than that of the simple networks. It is noteworthy to mention that the phenotypic variance of duplicated networks is on average significantly larger than that of simple networks with the same number of nodes, but not having the typical ‘duplicated’ network structure (Fig. 3*B*, comparing, for instance, the mean variances of the output nodes from the 20-node duplicated networks with those of simple 40-node networks (red arrows) (see also *SI Appendix*, Fig. S3.2). Similarly, the average phenotypic variance σ as measured by multiplying variance of both (mean) output node values of populations (1000) of simple networks of 10 nodes and their duplicated networks is more than four times higher for duplicated (*σ*=0.176) compared to single networks (*σ*=0.041; Fig. 3*C*). For each pair of simulated simple-duplicated networks, the phenotypic vector length increases with about 30% (average vector length for single networks: 0.77±0.31; for duplicated networks: 1.00±0.30). Furthermore, this increase in phenotypic value (trait) is in the same direction as the phenotype of the single network. The relative angle between both vector angles is 0°±5° (Fig. 3*D*). This holds true for networks of all sizes and initialization conditions (see *SI Appendix* S2 and S3). In other words, genome doubling affects the phenotypic trait in the same ‘direction’ as in the simple ‘ancestral’ network, but in a multiplicative manner. For instance, if we consider gene expression as a trait, when gene expression is at a certain ‘high level’ in a simple network, gene expression will generally be further increased in the duplicated network. The same is true for repression of gene expression: in the duplicated network, gene repression will be higher/stronger. Of course, there are exceptions to the rule, indicated by vectors that point in contradictory directions, e.g., vectors in the lower left quadrant that point upwards rather than downwards (Fig. 3*A*). Therefore, we show that the duplication of the particular structure of a genetic network, rather than the increase in nodes, underlies the observed pattern of phenotypic (or niche) expansion. WGD thus seems to systematically ‘exaggerate’ the obtained phenotypic value of the single network rather than driving it into a random direction.

Selection along environmental gradients. The relative fitness between the simple and duplicated networks, as for instance expressed in differential survival or reproduction, is higher for phenotypes generated by simple networks in the reference environment, and when deviations between the new and reference environment are small. In contrast, when environmental change is large(r), the fitness of the duplicated networks exceeds that of the simple networks (Fig. 4, *A* and *C*). Comparing relative fitness rates (***w***_*duplicated*_ –***w***_*single*_)/***w***_*single*_, it becomes obvious that even small mismatches between the environment and phenotype impose a strong selection against the duplicated genomes (Fig. 4, *B* and *D*). With increasing environmental change, the fitness of the duplicated networks exceeds that of the simple networks, indicating that they will be favored compared to the non-duplicated networks. Fitness differences decrease with an increased number of nodes in the single network. When environmental change is too large, fitness differences equalize at zero since none of the networks can persist. This pattern is not affected by the number of nodes in the single network.

**Fig. 4.**
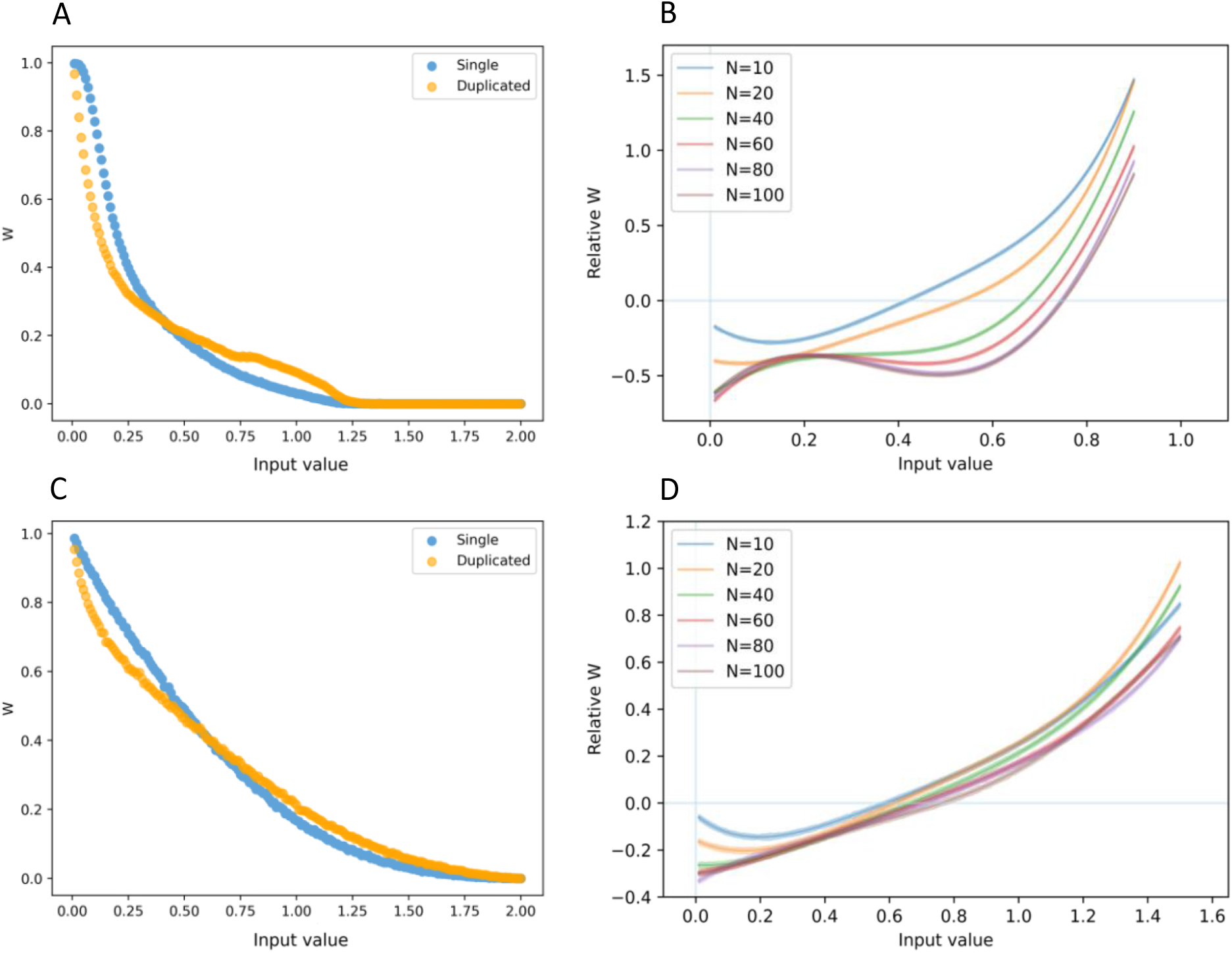
Fitness of simple and duplicated networks relative to the fitness of the population of simple networks in the reference environment, as a function of different input values, assuming a Gaussian fitness function (eq. 3) (A) and a linear fitness function (eq. 2) (C). Differences in survival rate as a function of different input values (assuming a Gaussian fitness function (eq. 3) (B) and a linear fitness function (eq. 2) (D) of input value, for networks of different size. Note that we only report relative w for input values leading to non-zero values for the single network.

## Discussion

The longer-term consequences of WGDs have been discussed at large. Whole genome duplications increase both mutational and environmental robustness due to redundancy and increased genetic variation (Van de Peer et al. 2009; te Beest et al. 2012; Van de Peer et al. 2017; Baduel et al. 2018; Doyle and Coate 2019; Van de Peer et al. 2021). Many studies have reported on the co-option of the extra duplicates specifically retained following WGD in different biological processes or pathways, increasing biological complexity or creating biological novelty (Sato et al. 2012; Cai et al. 2015; Unver et al. 2017; Zhang et al. 2020). However, rewiring of gene interactions and functional divergence of genes takes time and selective advantages of WGDs on the short – or even immediate – term, often remain elusive. We have previously wondered about the ‘conundrum’ between the many examples of recurrent polyploidy and the existence of many polyploids of recent origin, which seem to contrast with the evidence of relatively few polyploidy events that have been established on the long-term, certainly within the same evolutionary lineage (Van de Peer et al. 2017; Carretero-Paulet and Van de Peer 2020; Wong et al. 2020). The long-term fixation of polyploidy does not seem to occur randomly in space and time. One notable example is the biased distribution of ‘survived’ WGD events across independent plant lineages at the Cretaceous–Paleogene or K-Pg boundary, about 66 million years ago (Mya) (Vanneste et al. 2014b). Other ‘waves’ of WGDs may correlate with periods of global climatic change during the Paleocene–Eocene, ca. 56–54 mya (Cai et al. 2019), or the last glaciations (Novikova et al. 2018). The possible correlation between the ‘establishment’ of WGDs at times of environmental upheaval is interesting, but, although some interesting hypotheses have been put forward (Freeling 2017; Levin and Soltis 2018), remains to be explained. The fact that polyploids can survive drastically changing conditions or cataclysmic impacts while their diploid progenitors cannot, suggests a short-term, perhaps even immediate, evolutionary advantage for polyploids.

Some of the immediate consequences of polyploidy have been well described (Baduel et al. 2018; Doyle and Coate 2019; Bomblies 2020). One of the most consistent effects of WGD is an increase in cell size, but also physiological effects have been often observed. For instance, Chao and colleagues (2013) (Chao et al. 2013) demonstrated that *Arabidopsis thaliana* first generation autotetraploids have instantaneously enhanced salt tolerance compared to their diploid progenitors. Neo-autotetraploid *A. thaliana* lines were shown to experience a tradeoff, being less fit compared to diploid progenitors under non-saline conditions, but more fit in response to saline challenge. The authors proposed that in conditions of salinity stress the autopolyploid lineages would benefit from a fitness advantage that could contribute to their establishment and persistence. In turn, del Pozo and Ramirez-Parra showed that autotetraploid *A. thaliana* is also more drought tolerant (del Pozo and Ramirez-Parra 2014). Tetraploid rice (*Oryza sativa*) and citrange (*Citrus sinensis* L. Osb. x *Poncirus trifoliata* L. Raf.) too, have an increased tolerance to salt and drought stress because of WGD, which affects the expression of genes involved in stress and phytohormone response pathways (Yang et al. 2014; Ruiz et al. 2016). Similarly, tetraploid rootstock-grafted watermelon (*Citrullus lanatus*) plants are more tolerant to salt stress than are diploid plants (Zhu et al. 2018). Although such physiological and cellular responses to stress have thus been frequently documented for polyploids (Van de Peer et al. 2021), the exact molecular processes underlying these responses remain elusive (Fox et al. 2020). Both the ‘gigas’ effect shown by polyploids, as well as observations in shifts in photosynthetic rates or stress tolerance, are in line with our findings when considering polyploidy at the genomic and gene regulatory network level. As shown in Figure 3, simply considering the particular structure of duplicated networks, these networks show greater variation in trait values, solely likely being able to explain observations such as increased drought and salt tolerance.

However, in stable, non-changing environments, polyploidy will often be disadvantageous, as shown by our simulations, but also observed in vivo. For instance, in *Heuchera cylindrica*, an herbaceous perennial plant, increased nutrient requirements following polyploidy constrain the ability of new polyploids to establish in the nutrient-poor habitats the diploid progenitors thrive in (Anneberg and Segraves 2023). Similar observations were made for the autopolyploid complex *Dianthus broteri*, where, although higher ploidies have developed specific photochemical processes to survive in extremely warm conditions, the reduced performance of higher cytotypes render them less competitive in the ‘normal’ (non-stressed) environment (López-Jurado et al. 2020). Our simulation results suggest that duplicated networks – or their hosts - will be able to coexist to eventually replace their simple ancestors only under substantial environmental change (Fig. 4), for instance, when they end up in contrasting environments, or when the environment is quickly changing. These changes of the adaptive landscape might occur following major disturbances. During such events, any mutation must, by necessity, shift the value towards that new fitness peak if they are to increase fitness (Fig. 5). Or, in other words, when the original adaptive peak is sinking, overshooting is necessary to reach the new rising adaptive peak under discontinuous and/or fast environmental change (Fig. 5). Differential fitness is a first and foremost criterion underlying adaptive dynamics theory (Dercole and Rinaldi 2008).

We need to notice that, in contrast to our approach here, polyploids are continuously but in low frequencies produced by their non-duplicated ancestors. Polyploids emerge from meiotic failures that lead to unreduced gametes, which are documented to occur in low frequencies: 0.1 to 2% in vascular plants (Kreiner et al. 2017). This implies that every 1/1000 to 2/100 offspring (seeds) will experience this potential niche expansion. This number probably increases during times of environmental stress (De Storme et al. 2012; Van de Peer et al. 2021). Given high fecundity in most (WGD) plants, our assumption of a one-to-one doubling event is thus not too far from reality. This implies that any potential for establishment will depend on their fitness advantage compared to their ancestors and the level of standing genetic variation of the ancestor population. Since we show polyploids to have ‘exaggerated’ traits of their ancestors, fitness advantages are to be expected mainly when rapid and drastic environmental change is already in line with earlier ambient selection. For instance, when ancestor populations evolved under continuous warming, extreme heat waves will promote polyploid invasion. If such a period of warming would be followed by extreme cold, fitness disadvantages would disappear because the extreme phenotype would then exaggerate evolved maladaptations. On the other hand, we do observe - albeit much rarer - cases where the orientation of vectors in trait-space (Fig. 3, *A* and *D*), describing the relative contributions of the output traits to divergence between simple and duplicated networks, are almost opposite. In such cases, even when the fitness landscape changes more drastically, causing niche shifts rather than niche expansion, polyploids might be the ‘hopeful monsters’ being able to adapt, where their diploid progenitors go extinct (Fig. 5). This way, whole genome duplication or polyploidy might even explain large ‘jumps’ in evolution, or so-called saltational evolution (Theissen 2009). We thus show that immediate consequences of polyploidy can be significant, but it remains to be further tested whether they can indeed explain the preferential survival of polyploids over diploids during cataclysmic impacts that are generated by completely different selection pressure, and times of extinction, as previously suggested (Lohaus and Van de Peer 2016; Van de Peer et al. 2017; Carretero-Paulet and Van de Peer 2020). A more explicit eco-evolutionary approach that considers the history of selection, magnitude, frequency, and direction of environmental change is needed to provide the needed answers in these. This remains the topic of future work.

**Fig. 5.**
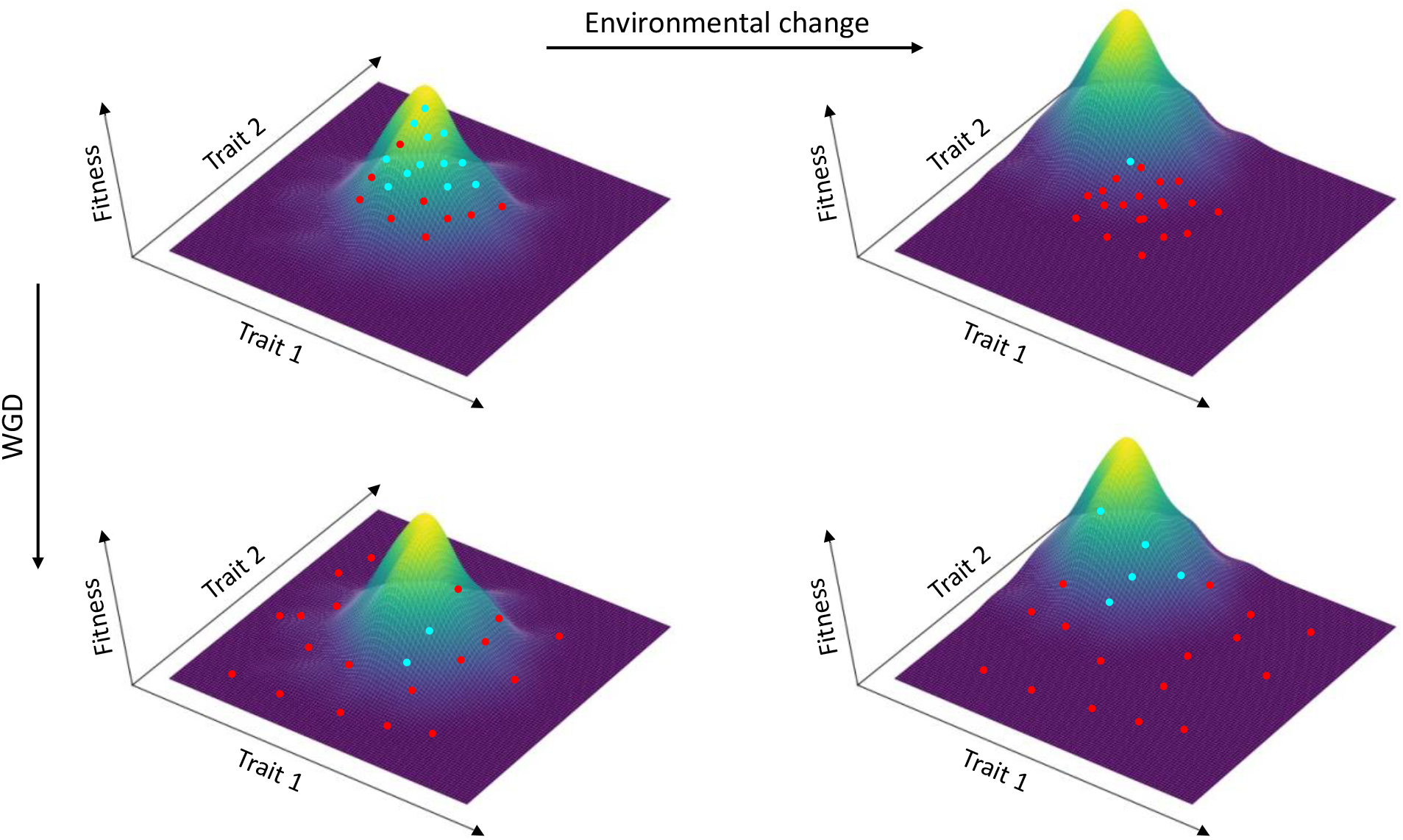
3D representation of fitness landscapes in which hills, corresponding to local adaptive peaks, are surrounded by valleys or depressions, corresponding to regions of the phenotype space where no survival is possible. Polyploidy may allow a wider and faster exploration of phenotypic space, ultimately conferring a potential adaptive advantage under challenging environmental conditions. Blue-green dots are individuals that can survive, red dots denote organisms that cannot survive. In a stable environment (top left panel), non-polyploid organisms are expected to have reached their local adaptive peaks. WGD results in an expansion of the phenotypic space covered by the population, although some polyploid genotypes might survive, most polyploids cannot survive in this environment (bottom-left panel). Adaptive landscapes are readily distorted by environmental challenges, such as cataclysmic or extinction events (right panels), resulting in shifts in the relative locations of their adaptive peaks. Under these conditions, although most diploids are expected to perish (top right panel), some polyploid organisms (which could be referred to as ‘hopeful monsters’), featured by wider accessible phenotype space (see text for details), have better chances to fall near the peak of a newly formed adaptive hill and thus to acquire the necessary evolutionary innovations to colonize novel niches (bottom right panel).

## Acknowledgements

We thank Yao Yao (Biocomputing and Developmental Systems Group, University of Limerick, Limerick, Ireland) and Felipe Kauai (VIB-Ghent University, Ghent, Belgium) for helpful discussions. This work was supported by the European Research Council (ERC) under the European Union’s Horizon 2020 research and innovation program (No. 833522) and from Ghent University (Methusalem funding, BOF.MET.2021.0005.01) (to Y.V.d.P.) and Ghent University (UGent BOF/STA/202009/039, BioGraph BOF.24Y.2019.0010.01) (to P.A.).

## Notes

### Competing Interest Statement

The authors have declared no competing interest.

## References

Adams DC, Collyer ML. 2009. A general framework for the analysis of phenotypic trajectories in evolutionary studies. Evolution 63: 1143–1154.

Albert R. 2005. Scale-free networks in cell biology. J Cell Sci 118: 4947–4957.

Anneberg TJ, Segraves KA. 2023. Neopolyploidy causes increased nutrient requirements and a shift in plant growth strategy in Heuchera cylindrica. Ecology doi:10.1002/ecy.4054:e4054.

Baduel P, Bray S, Vallejo-Marin M, Kolář F, Yant L. 2018. The “Polyploid Hop”: Shifting challenges and opportunities over the evolutionary lifespan of genome duplications. Front Ecol Evol 6: 117.

Barabasi AL. 2009. Scale-free networks: a decade and beyond. Science 325: 412–413.

Barabasi AL, Bonabeau E. 2003. Scale-free networks. Sci Am 288: 60–69.

Barker MS, Arrigo N, Baniaga AE, Li Z, Levin DA. 2016. On the relative abundance of autopolyploids and allopolyploids. New Phytologist 210: 391–398.

Bolnick DI, Barrett RDH, Oke KB, Rennison DJ, Stuart YE. 2018. (Non)Parallel evolution. Ann Rev Ecol Evol Syst 49: 303–330.

Bomblies K. 2020. When everything changes at once: finding a new normal after genome duplication. Proc Biol Sci 287: 20202154.

Broido AD, Clauset A. 2019. Scale-free networks are rare. Nat Commun 10: 1017.

Cai J, Liu X, Vanneste K, Proost S, Tsai WC, Liu KW, Chen LJ, He Y, Xu Q, Bian C et al. 2015. The genome sequence of the orchid Phalaenopsis equestris. Nat Genet 47: 65–72.

Cai L, Xi Z, Amorim AM, Sugumaran M, Rest JS, Liu L, Davis CC. 2019. Widespread ancient whole-genome duplications in Malpighiales coincide with Eocene global climatic upheaval. New Phytol 221: 565–576.

Carretero-Paulet L, Van de Peer Y. 2020. The evolutionary conundrum of whole genome duplication. Am J Bot 107: 1101–1105.

Chang J, Duong TA, Schoeman C, Ma X, Roodt D, Barker N, Li Z, Van de Peer Y, Mizrachi E. 2023. The genome of the king protea, Protea cynaroides. Plant J 113: 262–276.

Chao DY, Dilkes B, Luo H, Douglas A, Yakubova E, Lahner B, Salt DE. 2013. Polyploids exhibit higher potassium uptake and salinity tolerance in Arabidopsis. Science 341: 658–659.

Comai L. 2005. The advantages and disadvantages of being polyploid. Nat Rev Genet 6: 836–846.

Cooper MB, Loose M, Brookfield JF. 2008. Evolutionary modelling of feed forward loops in gene regulatory networks. Biosystems 91: 231–244.

Cuypers TD, Hogeweg P. 2014. A synergism between adaptive effects and evolvability drives whole genome duplication to fixation. PLoS Comput Biol 10: e1003547.

Cuypers TD, Rutten JP, Hogeweg P. 2017. Evolution of evolvability and phenotypic plasticity in virtual cells. BMC Evol Biol 17: 60.

De Lisle SP, Bolnick DI. 2020. A multivariate view of parallel evolution. Evolution 74: 1466–1481.

De Storme N, Copenhaver GP, Geelen D. 2012. Production of diploid male gametes in Arabidopsis by cold-induced destabilization of postmeiotic radial microtubule arrays. Plant Physiol 160: 1808–1826.

Dehal P, Boore JL. 2005. Two rounds of whole genome duplication in the ancestral vertebrate. PLoS Biol 3: e314.

del Pozo JC, Ramirez-Parra E. 2014. Deciphering the molecular bases for drought tolerance in Arabidopsis autotetraploids. Plant Cell Environ 37: 2722–2737.

Dercole F, Rinaldi S. 2008. Analysis of Evolutionary Processes: The Adaptive Dynamics Approach and Its Applications. Princeton University Press.

Doyle JJ, Coate JE. 2019. Polyploidy, the nucleotype, and novelty: the impact of genome doubling on the biology of the cell. Int J Plant Sci 180: 1–52.

Fawcett JA, Maere S, Van de Peer Y. 2009. Plants with double genomes might have had a better chance to survive the Cretaceous-Tertiary extinction event. Proc Natl Acad Sci U S A 106: 5737–5742.

Force A, Lynch M, Pickett FB, Amores A, Yan YL, Postlethwait J. 1999. Preservation of duplicate genes by complementary, degenerative mutations. Genetics 151: 1531–1545.

Fox D, Soltis DE, Soltis PS, Ashman T-L, Van de Peer Y. 2020. Polyploidy: a biological force from cells to ecosystems. Trends Cell Biology 30: 688–694.

Freeling M. 2017. Picking up the Ball at the K/Pg Boundary: The Distribution of Ancient Polyploidies in the Plant Phylogenetic Tree as a Spandrel of Asexuality with Occasional Sex. Plant Cell 29: 202–206.

Herrera C, Zufiria PJ. 2011. Generating scale-free networks with adjustable clustering coefficient via random walks. In 2011 IEEE Network Science Workshop, doi:10.1109/NSW.2011.6004642, pp. 167–172.

Koenen EJM, Ojeda DI, Bakker FT, Wieringa JJ, Kidner C, Hardy OJ, Pennington RT, Herendeen PS, Bruneau A, Hughes CE. 2020. The Origin of the Legumes is a Complex Paleopolyploid Phylogenomic Tangle Closely Associated with the Cretaceous–Paleogene (K–Pg) Mass Extinction Event. Syst Biol doi:10.1093/sysbio/syaa041.

Kreiner JM, Kron P, Husband BC. 2017. Evolutionary Dynamics of Unreduced Gametes. Trends Genet 33: 583–593.

Landis JB, Soltis DE, Soltis PS. 2017. Comparative transcriptomic analysis of the evolution and development of flower size in Saltugilia (Polemoniaceae). BMC Genomics 18: 475.

LeCun Y, Bengio Y, Hinton G. 2015. Deep learning. Nature 521: 436–444.

Leebens-Mack JH Barker MS Carpenter EJ Deyholos MK Gitzendanner MA Graham SW Grosse I Li Z Melkonian M Mirarab S et al. 2019. One thousand plant transcriptomes and the phylogenomics of green plants. Nature 574: 679–685.

Levin DA. 1975. Minority cytotype exclusion in local plant populations. Taxon 24: 35–43.

Levin DA, Soltis DE. 2018. Factors promoting polyploid persistence and diversification and limiting diploid speciation during the K–Pg interlude. Curr Opin Plant Biol 42: 1–7.

Lohaus R, Van de Peer Y. 2016. Of dups and dinos: evolution at the K/Pg boundary. Curr Opin Plant Biol 30: 62–69.

López-Jurado J, Balao F, Mateos-Naranjo E. 2020. Polyploidy-mediated divergent light-harvesting and photoprotection strategies under temperature stress in a Mediterranean carnation complex. Environ Exp Bot 171: 103956.

Moreira A, Rennó-Costa C. 2021. Evolutionary Strategies Applied to Artificial Gene Regulatory Networks. doi:10.1101/2021.09.28.462218. bioRxiv.

Morgan C, Zhang H, Henry CE, Franklin FCH, Bomblies K. 2020. Derived alleles of two axis proteins affect meiotic traits in autotetraploid Arabidopsis arenosa. Proc Natl Acad Sci U S A 17: 8980–8988.

Mottes F, Villa C, Osella M, Caselle M. 2021. The impact of whole genome duplications on the human gene regulatory networks. PLoS Comput Biol 17: e1009638.

Novikova PY, Hohmann N, Van de Peer Y. 2018. Polyploid Arabidopsis species originated around recent glaciation maxima. Curr Opin Plant Biol 42: 8–15.

Nwankpa C, Ijomah W, Gachagan A, Marshall S. 2018. Activation Functions: Comparison of trends in Practice and Research for Deep Learning. https://doiorg/1048550/arXiv181103378.

Prettejohn BJ, Berryman MJ, McDonnell MD. 2011. Methods for generating complex networks with selected structural properties for simulations: a review and tutorial for neuroscientists. Front Comput Neurosci 5: 11.

Ruiz M, Quinones A, Martinez-Alcantara B, Aleza P, Morillon R, Navarro L, Primo-Millo E, Martinez-Cuenca MR. 2016. Tetraploidy Enhances Boron-Excess Tolerance in Carrizo Citrange (Citrus sinensis L. Osb. x Poncirus trifoliata L. Raf.). Front Plant Sci 7: 701.

Sato S Tabata S Hirakawa H Asamizu E Shirasawa K Isobe S Kaneko T Nakamura Y Shibata D Aoki K et al. 2012. The tomato genome sequence provides insights into fleshy fruit evolution. Nature 485: 635–641.

Schluter D. 2001. Ecology and the origin of species. Trends Ecol Evol 16: 372–380.

Scott AL, Richmond PA, Dowell RD, Selmecki AM. 2017. The Influence of Polyploidy on the Evolution of Yeast Grown in a Sub-Optimal Carbon Source. Mol Biol Evol 34: 2690–2703.

Selmecki AM, Maruvka YE, Richmond PA, Guillet M, Shoresh N, Sorenson AL, D. S, Kishony R, Michor F, Dowell R et al. 2015. Polyploidy can drive rapid adaptation in yeast. Nature 519: 349–352.

Soltis PS, Liu X, Marchant DB, Visger CJ, Soltis DE. 2014. Polyploidy and novelty: Gottlieb’s legacy. Philos Trans R Soc Lond B Biol Sci 369.

Soltis PS, Soltis DE. 2009. The Role of Hybridization in Plant Speciation. Annu Rev Plant Biol 60: 561–588.

Taylor JS, Braasch I, Frickey T, Meyer A, Van de Peer Y. 2003. Genome duplication, a trait shared by 22000 species of ray-finned fish. Genome Res 13: 382–390.

Te Beest M, Le Roux JJ, Richardson DM, Brysting AK, Suda J, Kubesova M, Pysek P. 2012. The more the better? The role of polyploidy in facilitating plant invasions. Ann Bot 109: 19–45.

Theissen G. 2009. Saltational evolution: hopeful monsters are here to stay. Theory Biosci 128: 43–51.

Tran NH, Choi KP, Zhang L. 2013. Counting motifs in the human interactome. Nat Commun 4: 2241.

Unver T, Wu Z, Sterck L, Turktas M, Lohaus R, Li Z, Yang M, He L, Deng T, Escalante FJ et al. 2017. Genome of wild olive and the evolution of oil biosynthesis. Proc Natl Acad Sci U S A 114: E9413–E9422.

Van de Peer Y, Ashman TL, Soltis PS, Soltis DE. 2021. Polyploidy: an evolutionary and ecological force in stressful times. Plant Cell 33: 11–26.

Van de Peer Y, Maere S, Meyer A. 2009. The evolutionary significance of ancient genome duplications. Nat Rev Genet 10: 725–732.

Van de Peer Y, Mizrachi E, Marchal K. 2017. The evolutionary significance of polyploidy. Nat Rev Genet 18: 411–424.

Vanneste K, Baele G, Maere S, Van de Peer Y. 2014a. Analysis of 41 plant genomes supports a wave of successful genome duplications in association with the Cretaceous-Paleogene boundary. Genome Res 24: 1334–1347.

Vanneste K, Baele G, Maere S, Van de Peer Y. 2014b. Analysis of 41 plant genomes supports a wave of successful genome duplications in association with the Cretaceous–Paleogene boundary. Genome Res 24: 1334–1347.

Wendel JF. 2015. The wondrous cycles of polyploidy in plants. Am J Bot 102: 1753–1756.

Wong GK, Soltis DE, Leebens-Mack J, Wickett NJ, Barker MS, Van de Peer Y, Graham SW, Melkonian M. 2020. Sequencing and Analyzing the Transcriptomes of a Thousand Species Across the Tree of Life for Green Plants. Annu Rev Plant Biol 71: 1.1–1.25.

Wu S, Han B, Jiao Y. 2020. Genetic Contribution of Paleopolyploidy to Adaptive Evolution in Angiosperms. Mol Plant 13: 59–71.

Yang PM, Huang QC, Qin GY, Zhao SP, Zhou JG. 2014. Different drought-stress responses in photosynthesis and reactive oxygen metabolism between autotetraploid and diploid rice. Photosynthetica 52: 193–202.

Yao Y, Carretero-Paulet L, Van de Peer Y. 2019. Using digital organisms to study the evolutionary consequences of whole genome duplication and polyploidy. PLoS One 14: e0220257.

Yao Y, Marchal K, Van de Peer Y. 2014. Improving the adaptability of simulated evolutionary swarm robots in dynamically changing environments. PLoS One 9: e90695.

Yao Y, Storme V, Marchal K, Van de Peer Y. 2016. Emergent adaptive behaviour of GRN-controlled simulated robots in a changing environment. PeerJ 4: e2812.

Zhang L, Chen F, Zhang X, Li Z, Zhao Y, Lohaus R, Chang X, Dong W, Ho SYW, Liu X et al. 2020. The water lily genome and the early evolution of flowering plants. Nature 577: 79–84.

Zhu H, Zhao S, Lu X, He N, Gao L, Dou J, Bie Z, Liu W. 2018. Genome duplication improves the resistance of watermelon root to salt stress. Plant Physiol Biochem 133: 11–21.

