## Supplementary Material for "The duplication of genomes and gene regulatory networks and its potential for evolutionary adaptation and survival"

**This PDF file includes:**

Supporting text

Figures S1.1 to S3.3 (9)

Tables S1.1 to S3.3. (8)

SI References

### Supplementary 1: Sensitivity analysis of the network output according to the output constraining function

The “Tanh” function is one commonly used activation function in the area of neural networks (1, 2). We used this constrained function as a baseline as it represents biological functions with phenotypes being constrained by developmental, physiological limits (since “tanh” function has the max and min of ‘+1’ and ‘-1’) and it is a suitable function to model regulatory relations either repressing or enhancing. All results presented in the main document result from the use of this tanh-function. We tested sensitivity of the outcome to this assumption by running a non-limited function *F(x) = x.* We here present the average differences for this function in phenotypic variances (variance on the mean from 1000 simulations) between the duplicated and single networks according to the number of nodes in the single network (Table S.1). In all these simulations, input values are sampled from U[-1,+1] and all weights are generated from standard normal distributions. As expected, unconstrained connector functions generate larger variances compared to the constrained Tanh function. As for the baseline model, variances increase massively with GNR doubling, and is independent of the initialization procedure (Table S1.2). Also relative angles of the vectors show the same pattern as for the tanh-function (Table S1.3-S.4).

Table S1.1: Differences in phenotypic variation and its standard deviation in vector lengths (phenotypic values) as defined between the duplicated and single networks according to the size of single network, under the unconstrained output function.

| Size of the single network | Mean differences of the phenotypic variation | SD of the differences in phenotypic variation |
|---|---|---|
| 10 | 5.27 | 25.8 |
| 20 | 21.1 | 59 |
| 40 | 89.3 | 276.3 |
| 60 | 98.2 | 241.2 |
| 80 | 222 | 839.5 |
| 100 | 122 | 368.7 |

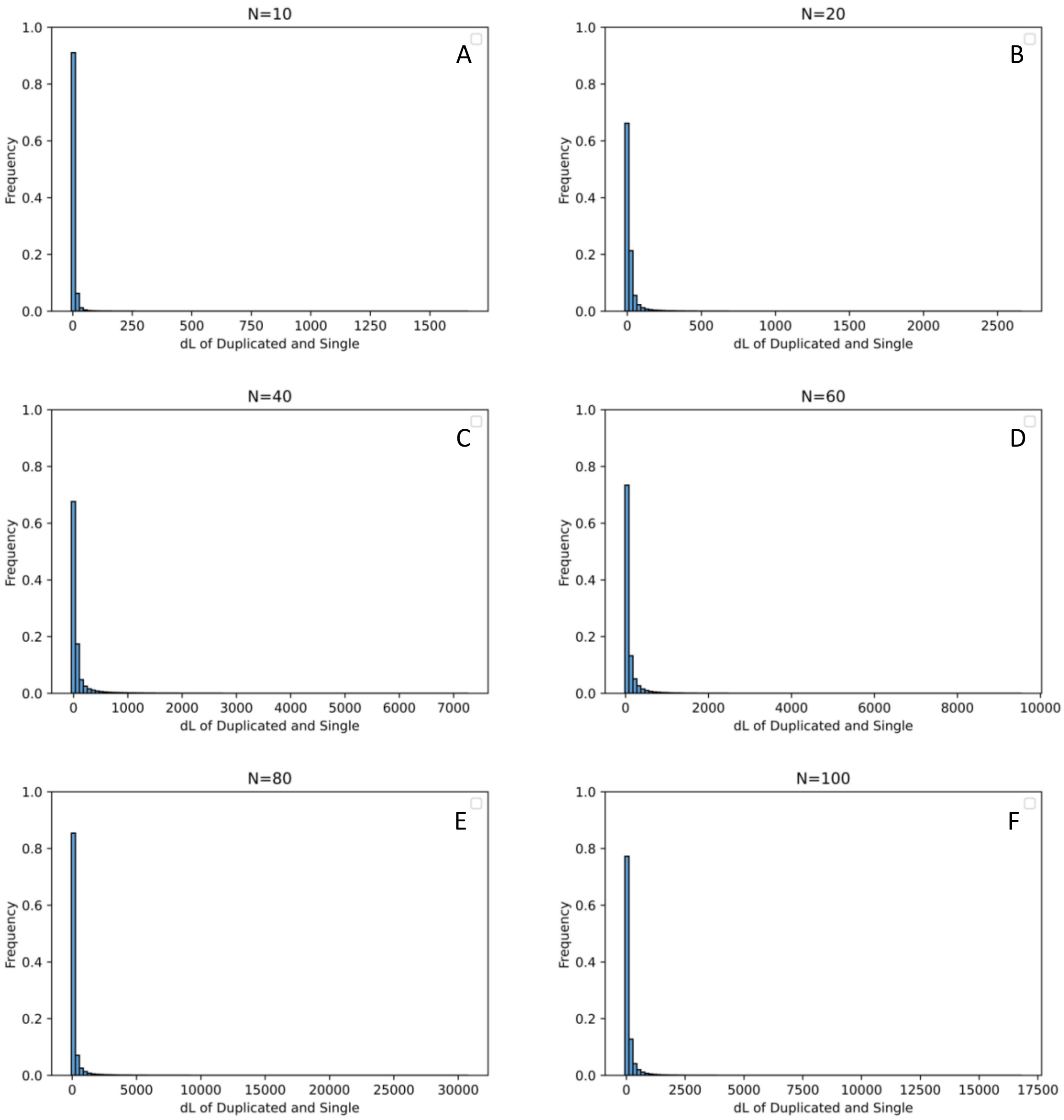


Figure S1.1. Distribution of differences in vector lengths (dL) between duplicated and single networks under the unconstrained function

Table S1.2. Differences in phenotypic variation and standard deviation between the duplicated and single networks of 10 nodes according to the used initialization (W: weight initialization, I: input initialization; N=Gaussian distribution N(0,1), U: Uniform distribution [-1,1]) under the unconstrained output function

| Scenario | Mean differences of the phenotypic variance | SD of the differences in phenotypic variance |
|---|---|---|
| W=N, I=N | 5.54 | 21 |
| W=U, I=U | 0.19 | 0.19 |
| W=N, I=U | 0.25 | 0.21 |
| W=U, I=N | 0.20 | 0.19 |

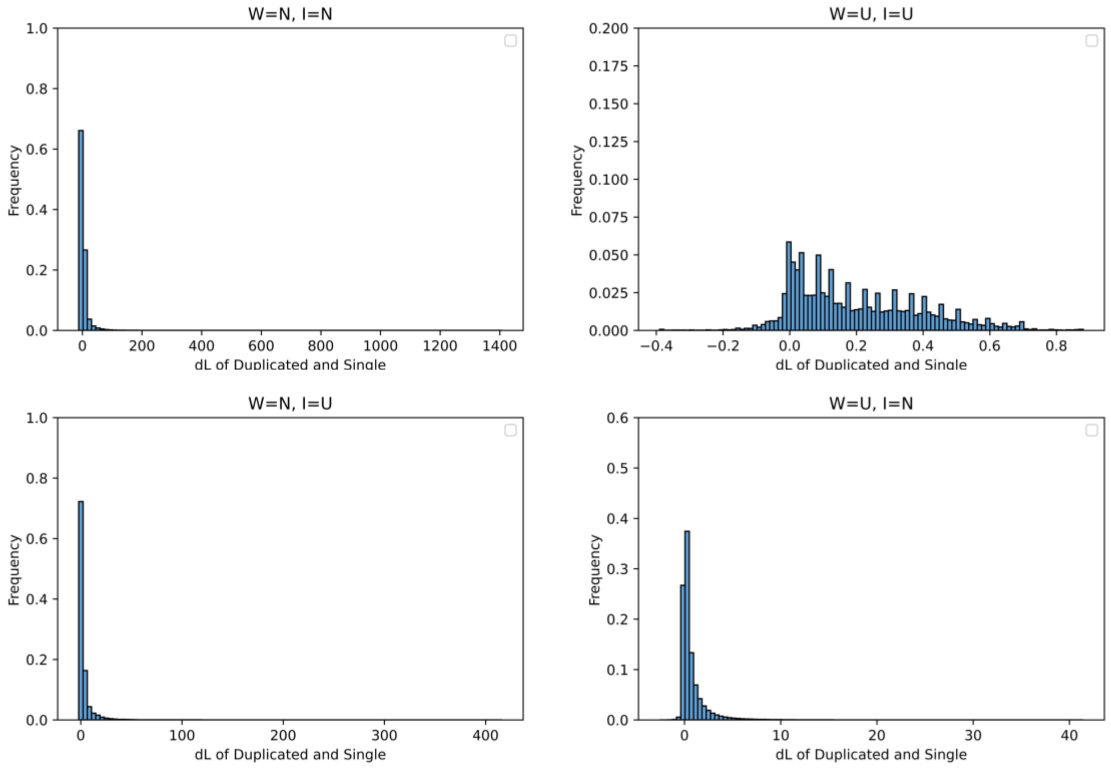


Figure S1.2: Distribution of differences in vector lengths (dL) between duplicated and single networks of 10 nodes according to the used initialization (W: weight initialization, I: input initialization; N=Gaussian distribution N(0,1), U: Uniform distribution [-1,1]) under the unconstrained output function.

Table S1.3: Relative angles and standard deviation between the phenotypic vectors of the duplicated and single networks, under the unconstrained output function for single networks of different size. All median values=0°.

| Size of the single network | Mean angle (α°) | SD angle (α°) |
|---|---|---|
| 10 | 3 | 38 |
| 20 | -4 | 52 |
| 40 | -1 | 55 |
| 60 | 4 | 54 |
| 80 | 11 | 68 |
| 100 | 0.43 | 80 |

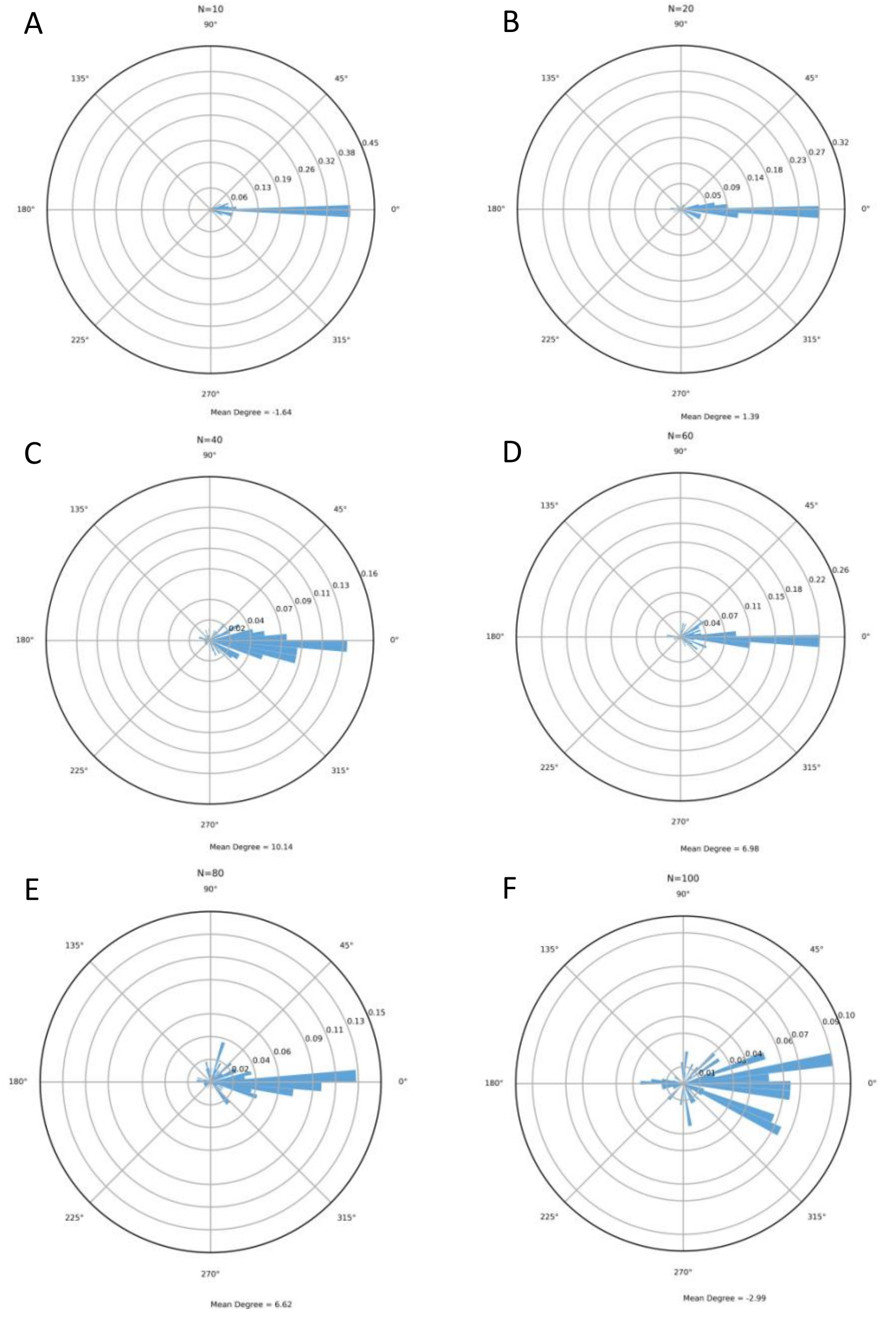


Figure S1.3: Distribution of the relative angles between the phenotypic vectors of the duplicated and single networks, under the unconstrained output function for single networks of different size.

Table S1.4: Relative angles and their standard deviation between the phenotypic vectors of the duplicated and single networks, under the unconstrained output function for the different initialization scenarios. (W: weight initialization, I: input initialization; N=Gaussian distribution N(0,1), U: Uniform distribution [-1,1]). All median values=0°.

| Scenario | Mean angle (α°) | SD angle (α°) |
|---|---|---|
| W=N, I=N | 3 | 45 |
| W=U, I=U | 5 | 28 |
| W=U, I=N | -4 | 42 |
| W=N, I=U | 1 | 31 |

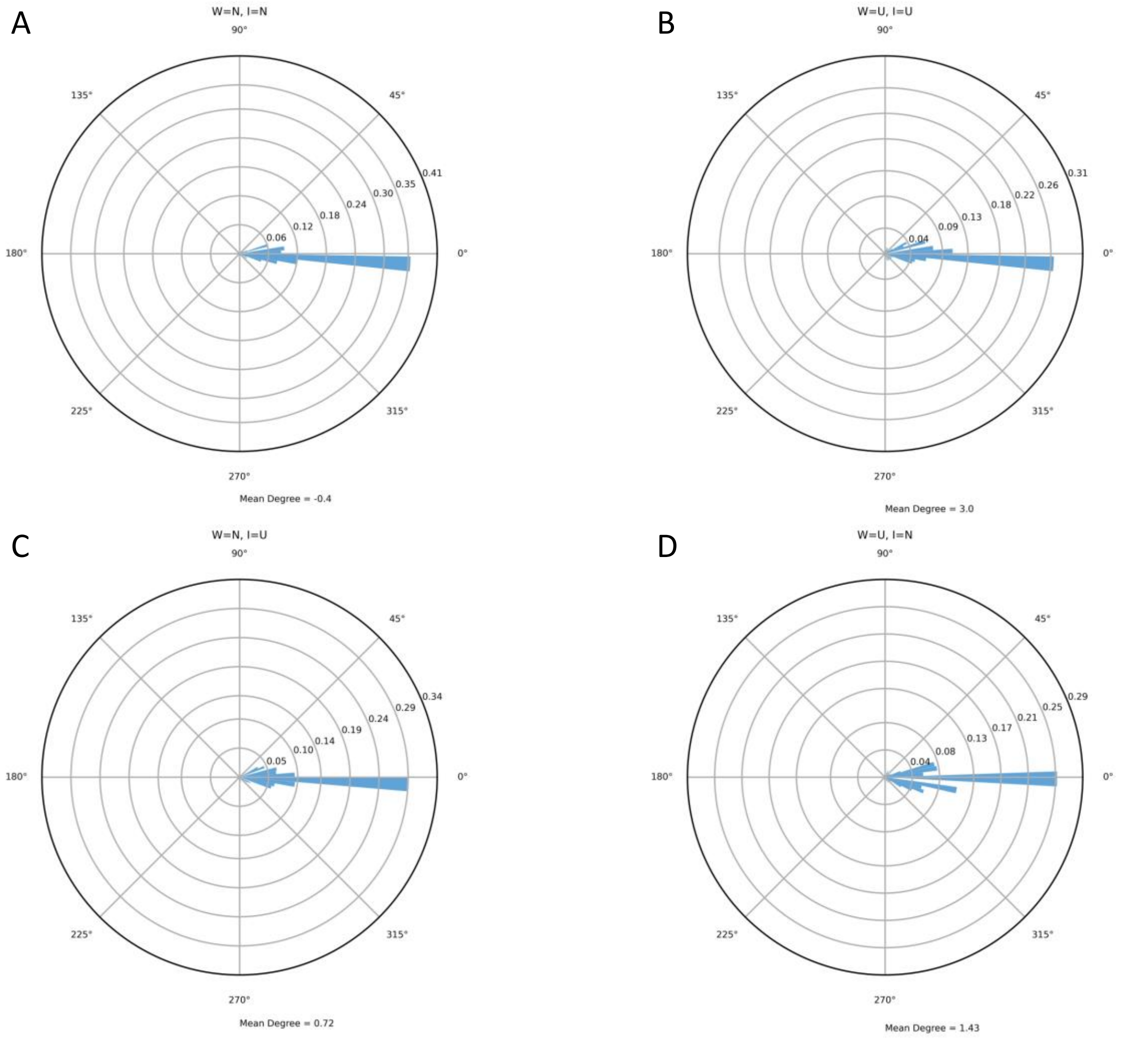


Figure S1.4: Distribution of relative angles between the phenotypic vectors of the doubled and single networks, under the unconstrained output function for the different initialization scenarios. (W: weight initialization, I: input initialization; N=Gaussian distribution N(0,1), U: Uniform distribution [-1,1]).

### Supplementary 2: Sensitivity on the network output according to the number of network nodes

In the main manuscript, we show that the variance is increasing with node number, and that duplication is having significant impact on the variance increase (Fig 3.2). A more profound analysis on the phenotypic changes from each paired single and duplicated network shows that changes in phenotypic vector length between the bivariate output nodes of the single and its duplicated network are qualitatively similar across the scenarios where single networks are built with different node numbers (Table S2.1).

Table S2.1. Differences in phenotypic vector lengths and their standard deviation as defined between the doubled and single networks according to the size of single network, under the Tanh function.

| Size of the single network | Mean differences of the phenotypic variation (vector length) | SD of the differences in phenotypic variation (vector length) |
|---|---|---|
| 10 | 0.25 | 0.22 |
| 20 | 0.33 | 0.25 |
| 40 | 0.34 | 0.27 |
| 60 | 0.31 | 0.29 |
| 80 | 0.28 | 0.32 |
| 100 | 0.24 | 0.32 |

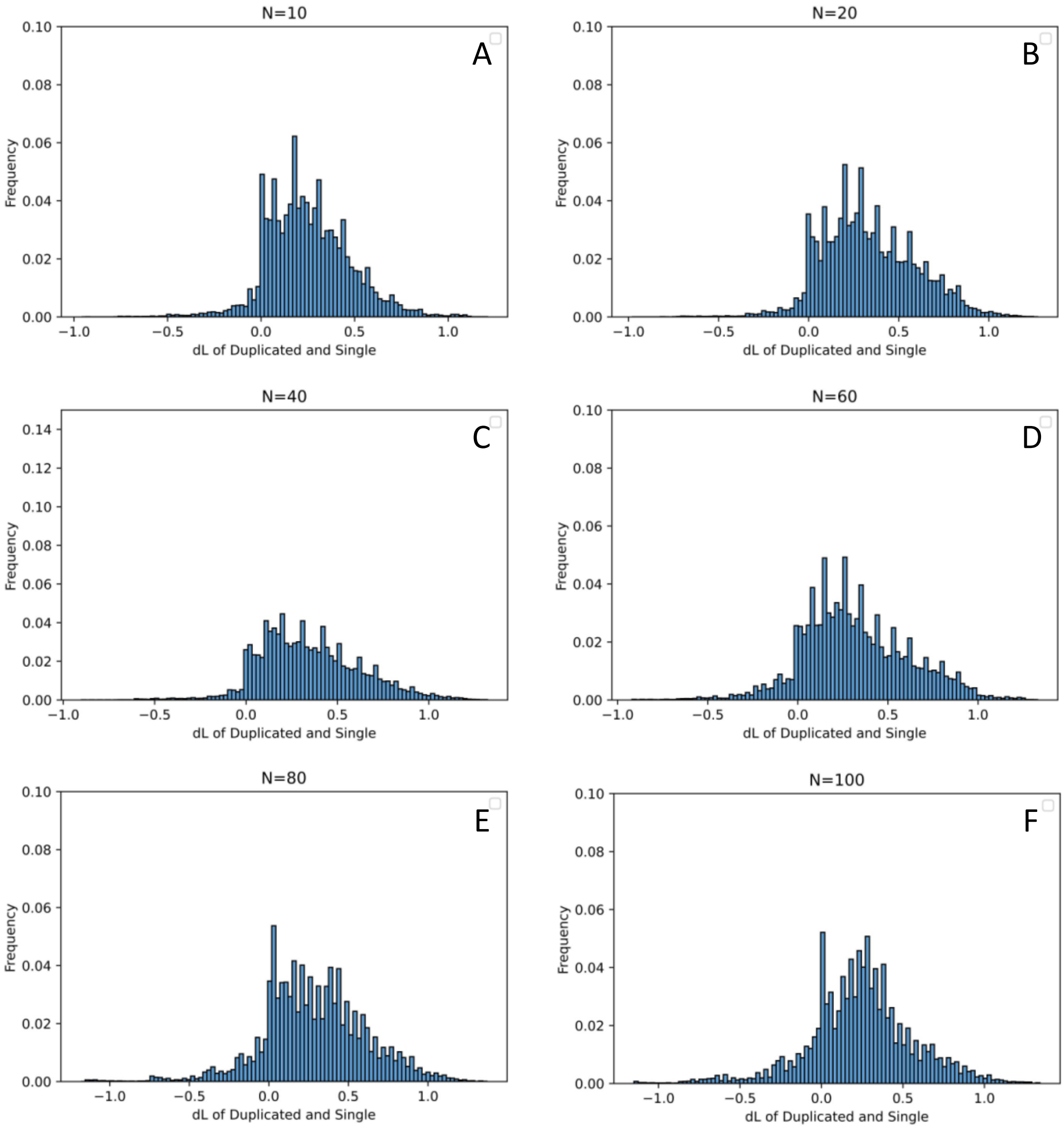


Figure S2.1: Distribution of differences in phenotypic vector lengths as defined between the doubled and single networks according to the size of single network, under the Tanh function.

Table S2.2: Relative angles and standard deviation between the phenotypic vectors of the doubled and single networks according to the size of the single network, under the Tanh function. All median values=0°.

| Size of the single network | Mean angle (α°) | SD angle (α°) |
|---|---|---|
| 10 | -2 | 27 |
| 20 | 0.54 | 41 |
| 40 | 5 | 43 |
| 60 | 0.05 | 54 |
| 80 | -2 | 64 |
| 100 | 5 | 70 |

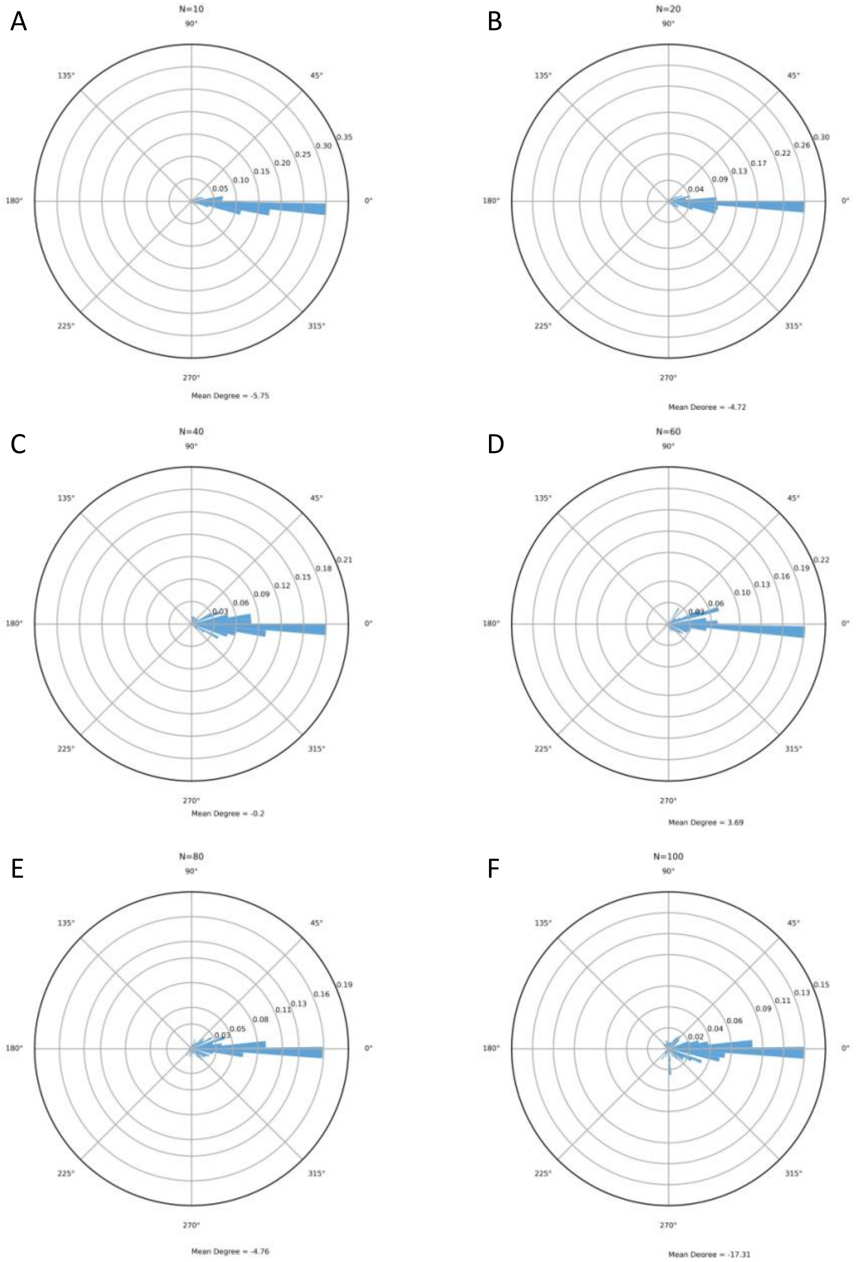


Figure S2.2: Distribution of relative angles and standard deviation between the phenotypic vectors of the doubled and single networks according to the size of the single network, under the Tanh function.

### Supplementary 3: Sensitivity analysis of the network output according to the input and edge weight distributions

Input values of the aGRNs for the results presented in the main manuscript were generated from a standard normal distribution N(0, 1). We tested for alternative input value distributions: (a) both edge weights and inputs sampled from a N(0, 1), (b) input values also sampled from a uniform random generator between -1 and +1 (U[-1,+1]) (beside weights of the edges), and (c) input generated from a U[-1,+1] and edges from a Gaussian N(0,1). We generated 1000 different network topologies, and from each of these 10K output values [$x_o$,$y_o$] were generated from 10K doubles of input values [$x_i$,$y_i$]. The output values then generate the phenotypic space based on 1000 values in the 2D plane.

We calculated differences in the phenotypic output values in several ways. First, the phenotypic space as determined by the mean of the two output values, was compared between the 1000 generated single and duplicated networks. As evidenced from Figures S3.1 and Table 3.1 differences were on average positive, indicating that our finding of increased variance after WGD remains conserved with different input and edge weight initialization distributions.

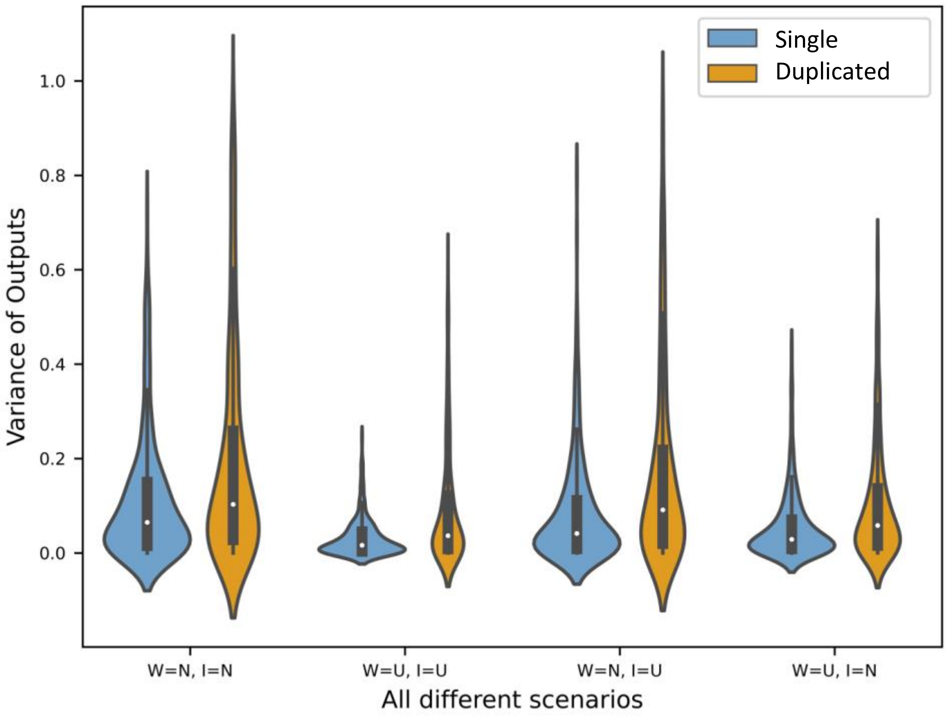


Figure S3.1 Violin plot of the differences in variances between doubles and single networks according to the values distributions of the network initialization. W=weight values, I= input values; N implies sampling from a standard Gaussian distribution N(0, 1); U implies sampling from a U[-1,+1]. Left panel the raw values distribution, right panel the mean and quartile distribution.

Table S3.1. Differences in phenotypic variance and its standard deviation between the duplicated and single networks according to the used initialization (W: weight initialization, I: input initialization; N=Gaussian distribution N(0,1), U: Uniform distribution [-1,1]) under the Tanh output function.

| Scenario | Mean differences of the phenotypic variance | SD of the differences in phenotypic variance |
|---|---|---|
| W=N, I=N | 0.22 | 0.21 |
| W=U, I=U | 0.19 | 0.19 |
| W=N, I=U | 0.25 | 0.21 |
| W=U, I=N | 0.20 | 0.19 |

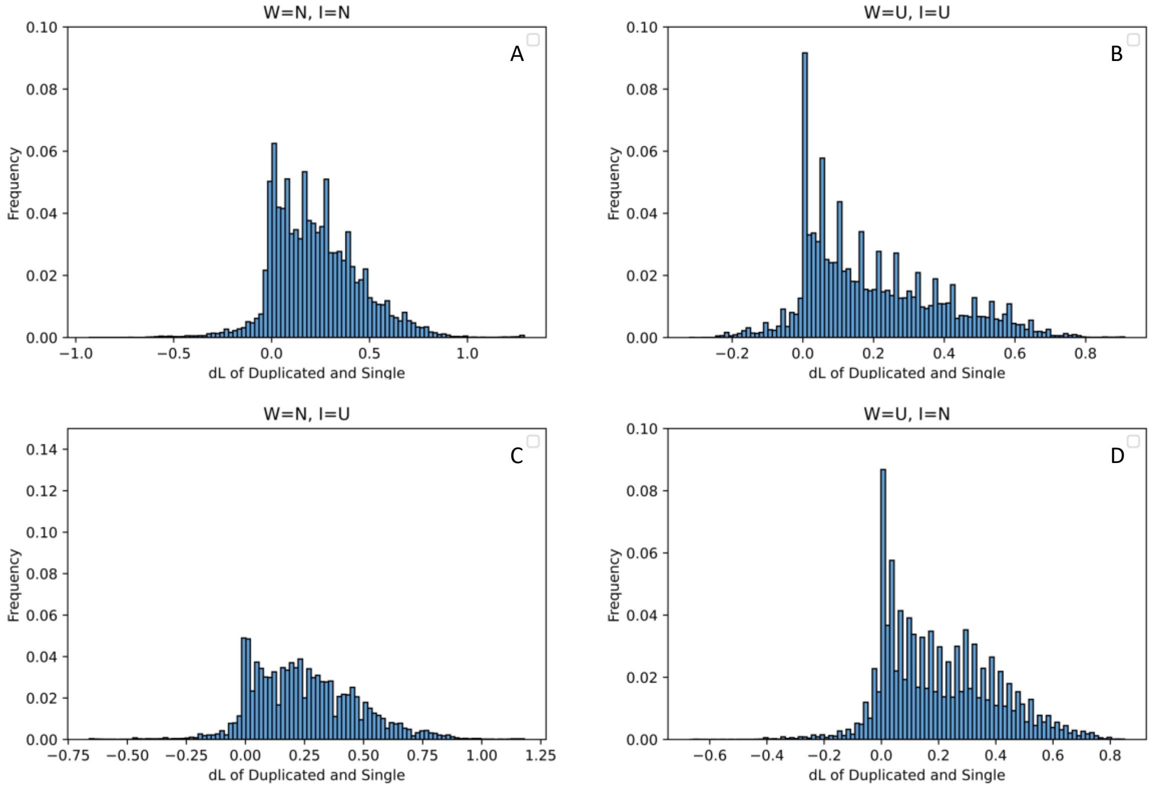


Figure S3.2: Distribution of differences in phenotypic vector length between the duplicated and single networks according to the used initialization (W: weight initialization, I: input initialization; N=Gaussian distribution N(0,1), U: Uniform distribution [-1,1]) under the Tanh output function.

A more profound analysis on the phenotypic changes from each paired single and duplicated network confirms the generality of our findings that the net differences in phenotypic value change as determined by the vector length in the 2D phenotypic space is positive, hence indicating that doubling on average increases the phenotypic value compared to those from the simple network (Fig. S3.2). Only when both the input and edge values were generated from a Gaussian distribution, a higher frequency of negative changes in vector length (10.1% negative rate for W=N I=N, 7.7% for W=N I=U, and 8.9% for W=U I=N, and 9.2% for W=U I=U), and hence shrinking of the phenotypic space, was observed.

Angular dispersion of the phenotypic vectors are equally centered around zero for all scenarios (Table 3.3 & Fig. S3.3), indicating that the initialization procedure neither impact the angular correlation of the phenotypic vectors after doubling. In all scenarios, the correlation between the phenotypic vector angles from the single and doubled networks was distributed around zero.

Table S3.3. Relative angles and standard deviation between the phenotypic vectors of the doubled and single networks, under the unconstrained output function for the different initialization scenarios. (W: weight initialization, I: input initialization; N=Gaussian distribution N(0,1), U: Uniform distribution [-1,1]). All median values=0°.

| Scenario | Mean angle (α°) | SD angle (α°) |
|---|---|---|
| W=N, I=N | -1 | 23 |
| W=U, I=U | 3 | 35 |
| W=U, I=N | -3 | 35 |
| W=N, I=U | -1 | 34 |

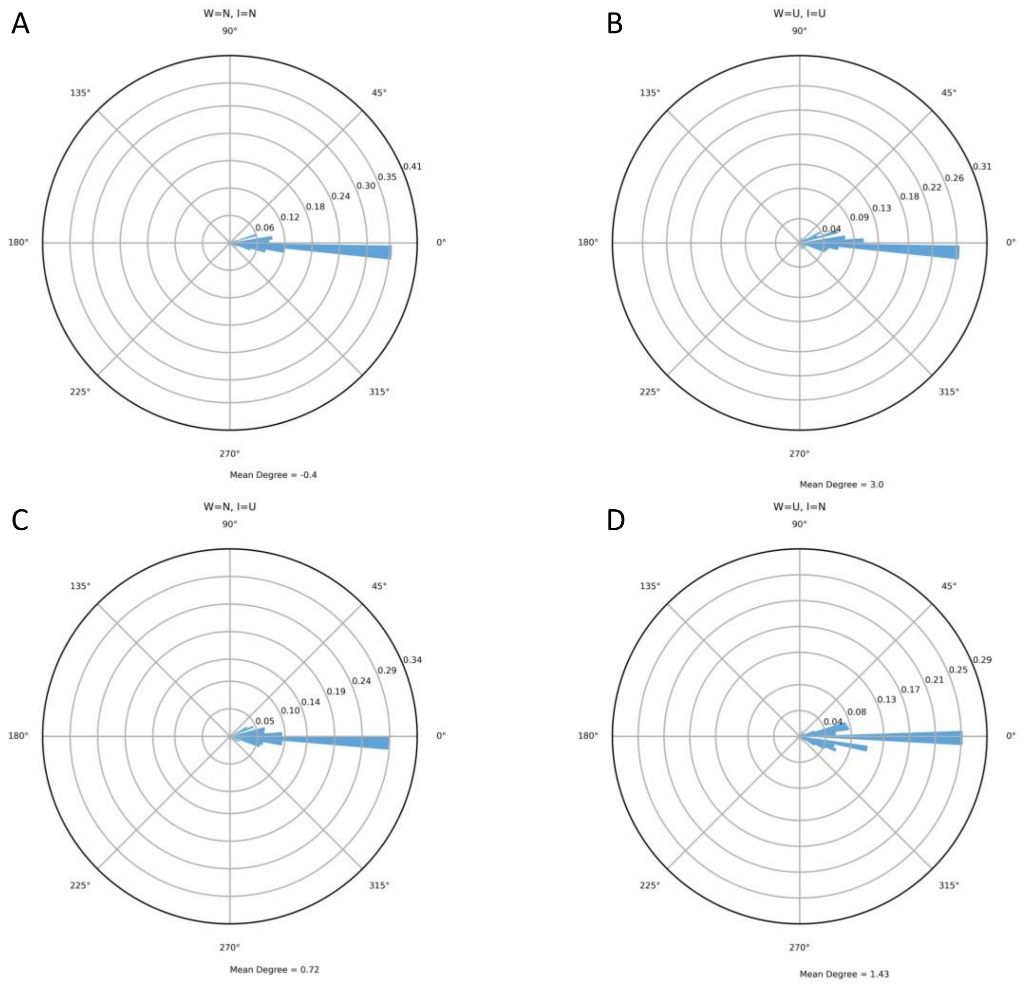


Figure S3.3. Distribution of relative angles between the phenotypic vectors of the doubled and single networks, under the unconstrained output function for the different initialization scenarios. (W: weight initialization, I: input initialization; N=Gaussian distribution N(0,1), U: Uniform distribution [-1,1]).

**SI References**


1. Y. LeCun, Y. Bengio, G. Hinton, Deep learning. *Nature* **521**, 436-444 (2015).
2. A. Moreira, C. Rennó-Costa (2021) Evolutionary strategies applied to artificial gene regulatory networks. *bioRxiv* 2021.09.28.462218.
